# Pre-treatment with the methanol extract of *Withania somnifera* prevents Diazinon-induced cardiotoxic effects of organophosphate poisoning

**DOI:** 10.1101/042523

**Authors:** Irungu Eric Mwangi, Mwangi Peter Waweru, Bukachi Frederick

## Abstract

Organophosphate poisoning represents a major and growing global health problem especially in the developing countries and cardiotoxicity is the major cause of death. Thus, a compelling need to develop novel low cost efficacious agents to manage this condition.

**Objective:** To evaluate the methanol extract of *Withania somnifera* as a pre-treatment agent in the prevention of the cardiotoxic effects of diazinon in Sprague Dawley rats

**Materials and Methods:** Twenty one (21) adult rats were randomized to receive 200 mg/kg methanol extract of *Withania somnifera* (test group), vehicle (negative control) or 200 μg/kg Neostigmine as pre-treatment 30 minutes prior to the oral administration of 200 mg/kg Diazinon. Baseline and post-treatment electrocardiograms (ECGs) were recorded by the Powerlab data acquisition system (ML865 AD instruments, Sydney, Australia). The experimental data were expressed as median ± the inter-quartile range and analysed using the Kruskal – Wallis non-parametric test and followed by Mann–Whitney U post hoc test in cases of significance, which was set at p < 0.05. Statistical Package for Social Sciences (SPSS) version 17 software was used for analysis.

**Results:** Pre-treatment with the methanol extract of *W. somnifera* had significant effect on the following diazinon-induced electrocardiographic changes; RR interval (0.026 (0.007 – 0.065) vs. 0.035 (0.019 – 0.050) vs. 0.090 (0.071 – 0.01), p = 0.031),heart rate (-54.235 (-115.317 – (-19.857)) vs. -−96.136 (-96.472 – (-43.879)) vs. -−174.361 (-189.775 – (-129.469)), p = 0.014), PR interval (0.006 (0.004 – 0.008) vs. 0.003 (0.001 – 0.004) vs. 0.009 (0.006 – 0.015), p = 0.019), QRS interval (0.005 (0.001 – 0.008) vs. -−0.002 (-0.005 – 0.001) vs. 0.007 (0.003 – 0.011), p = 0.023) and ST height (-34.830 (-63.578 – 4.215) vs. -−22.330 (-38.383– (-4.159)) vs. -−73.156 (-214.022– (-52.449)), p = 0.023). It however had no significant effect on the QTc interval changes (-0.005 (-0.011 – 0.003) vs. -−0.005 (-0.015 – 0.065) vs. -−0.021 (-0.060– (-0.006)), p = 0.174).

**Conclusion:** The efficacy of pre-treatment with the methanol extract of *Withania somnifera* was comparable to that of pre-treatment with Neostigmine a commonly used carbamate drug. Thus, it is a potentially viable low cost treatment option for organophosphate poisoning in resource-limited settings.

## 1.0. Introduction

It is estimated that over 80% of the population in developing countries rely partially or entirely on traditional systems of medicine (WHO, 2003). Organophosphate poisoning is a major global health problem (Eddleston et al., 2002) whose incidence is rising especially in the developing world. This is due to the introduction of large scale intensive agricultural practices which are often associated with the use of organophosphates as pesticides (Howard & Pope, 2002). This problem is often compounded by lack of basic training on safety practices in pesticide usage among most farm workers (Naidoo et al., 2010). The development of pesticide resistance often results in the use of higher doses of the chemicals further increasing risk of poisoning (Naidoo et al., 2010).

Organophosphate poisoning leads to cardiotoxicity that occurs early in poisoning leading to arrhythmogenesis, conduction abnormalities and myocardial ischemia and is the main cause of death (Bar-Meir et al., 2007; Saadeh et al., 1997).

The management of organophosphate poisoning whether by the use of pre-treatment or antidotal approaches after pesticide exposure remains unsatisfactory (Buckley, 2005). Attention has therefore turned to the investigation of compounds of herbal origin as putative sources of novel and efficacious remedies of organophosphate poisoning (Wang et al., 2011). Galanthamine from *Leucojum aestivum* and Huperzin A from *Huperiza serrate* are salutary examples and both have been evaluated as pre-treatment agents in organophosphate poisoning (van Helden et al., 2011).

Our study aimed to evaluate the effect of pre-treatment with the methanolic extract of *Withania somnifera* on the electrocardiogram (ECG) in Sprague Dawley rats exposed to toxic doses of diazinon, a commonly used organophosphate pesticide. This herb has multiple indications in Aryuvedic medicine (Singh and Kumar, 1998)

## 2.0. Methods

### 2.1. Preparation of extract

The plant material was collected on the outskirts of Athi River town located 30 kilometres from Nairobi, Kenya. The identity of the collected plant material was verified by plant taxonomists at the herbarium, School of Biological Sciences, University of Nairobi (Voucher number 100412).

The whole plant was air dried for one week. Thereafter, the material was milled to a fine powder using a standard kitchen blender. The extract was prepared by macerating the plant material with laboratory grade pure methanol in a 1:10 weight volume ratio for six hours. The solvent was then removed via vacuum evaporation in a rotary evaporator (Ugo Basile^®^) at 40°C and 376 Pascals pressure. The extract was then weighed and placed in an airtight sample bottle and refrigerated at 4°C.

### 2.2. Experimental Animals

Twenty one (21) adult (12 – 16 weeks old) male albino (Sprague-Dawley) rats weighing 250 g ± 50 g were selected for the study. They were kept at room temperature (~25°C) in a controlled light / dark environmental cycle of 9/15 hours. The animals were acquired locally and were housed in spacious cages at the animal house within the department of Medical Physiology, University of Nairobi, Chiromo Campus. They were provided with standard rat chow (Unga Feeds, Nairobi) and water *ad libitum*. There was a habituation period of one week prior to commencement of the experiments. All the experimental procedures used in the study were in compliance with the NIH Guide for the Care and Use of Laboratory Animals (2011).

### 2.3. Data collection

The experimental animals were randomized into three groups; the Positive Control, Test, and Negative Control groups with each group having seven animals (Table 1). Animals in the positive control group received Neostigmine (Tablets Limited, India) (200 μg/kg) while those in the test group received the extract (200 mg/kg). Those in the negative control group received 1 ml of vehicle (0.1 ml Dimethyl sulphoxide + 0.9 ml of Normal saline) as pre-treatment agent thirty (30) minutes before the administration of Diazinon (Sinochem Ningbo Chemicals Co. Ltd, China). The pre-treatment agents were administered via the intraperitoneal route. Diazinon was administered orally using a gavage needle. The dose of Diazinon used was 200 mg/kg, which was equivalent to 0.25 of the LD_50_ dose of Diazinon. Atropine sulphate (CSPC Group Company Limited, China) was administered intraperitoneally at a dose of 10 mg/kg ten (10) minutes after the administration of Diazinon to the experimental and positive control group animals. All the solutions administered to the animals were prepared just before administration.

**Table 1:** Table showing the Experimental Groups

| Positive Control | Test Group | Negative Control |
| --- | --- | --- |
| **NEOSTIGMINE**<br>+<br>**DIAZINON**<br>+<br>**ATROPINE** | **W. SOMNIFERA**<br>+<br>**DIAZINON**<br>+<br>**ATROPINE** | **DMSO**<br>+<br>**DIAZINON** |

#### 2.3.1. ECG Recording

Briefly, the ECG recording procedure was as follows; the rats were sedated using an intramuscular injection of Ketamine hydrochloride (Rotexmedica, Trittau, Germany) at a dose of 0.1 mg/g body weight. SKINTACT^®^ electrodes (Code T601, Leonhard Lang GmbH, Innsbruck, Austria) consisting of gel coated contact pads were placed on all the front and the left hind paws. Lead II ECGs were recorded using the Power lab data acquisition apparatus (ML865 AD instruments, Sydney, Australia). Default settings for the rat ECG were used for both the acquisition and the processing of the data. The ECGs were recorded at a range of two millivolts, a low pass filter of 50 Hertz and a high pass filter of 10 Hertz. Auto analysis of the data was carried out and verified manually. The following variables were analyzed:

Heart rate (HR) and RR interval, PR interval, QRS duration, ST segmentheight and QTc interval. The QTc interval was derived from the QT interval using Bazzet’s formula:

QTc=QT÷√RR interval.

The experimental data were presented as the median ± the inter-quartile range (Inter Quartile Range = Q1 – Q2 where Q1 is the 25th percentile and Q2 is the 75th percentile).

### 2.4. Statistical Analysis

The change in ECG measurements were analysed as the difference between the two readings ([post test] – [baseline]) for each parameter. The experimental data were analyzed using the Kruskal – Wallis non-parametric test and followed by Mann–Whitney U post hoc test in cases of significance, which was set at p < 0.05. Statistical Package for Social Sciences (SPSS) version 17 software was used for analysis.

## 3.0. Results

The percentage yield of the methanol extract was 1.67% of the dry weight of the powder.

All the rats survived until the end of the experiment. Sinus rhythm was present in all the baseline ECG recordings with the P wave preceding the QRS complexes. There was no significant difference in the baseline ECG parameters between the groups (Table 2). There was also no statistically significant difference in the change in QTc between the experimental groups (P = 0.174) (Figure 1). The other ECG parameters measured however showed significant differences as outlined below.

**Table 2:** Table showing median and inter quartile ranges of baseline ECG parameters

| ECG<br>Parameters | Group I<br>(Neostigmine)<br>(n=7) | Group II<br>(WS)<sup>‡</sup><br>(n=7) | Group III<br>(Diazinon)<br>(n=7) | P Value* |
| --- | --- | --- | --- | --- |
|  | Median<br>(IQR = Q1 – Q3)<sup>†</sup> |  |  |  |
| RR Interval<br>(s) | 0.142<br>(0.137 – 0.153) | 0.145<br>(0.138 – 0.153) | 0.143<br>(0.13 – 0.148) | 0.948 |
| HR<br>(b/min) | 423.172<br>(391.994 – 435.685) | 414.538<br>(390.611 – 440.116) | 432.416<br>(404.608 – 465.009) | 0.392 |
| PR Interval<br>(s) | 0.048<br>( 0.046 – 0.05) | 0.051<br>(0.043 – 0.053) | 0.044<br>(0.042- 0.049) | 0.221 |
| QRS Interval<br>(s) | 0.023<br>(0.02 – 0.03) | 0.028<br>(0.023 – 0.03) | 0.024<br>(0.023 – 0.028) | 0.400 |
| ST Height<br>(µV) | -6.636<br>(-57.64 – 1.333) | -27.844<br>(-51.262 - -14.433) | -43.369<br>(-93.748 - -2.489) | 0.749 |
| QTc (s) | 0.128<br>(0.125 – 0.190) | 0.127<br>(0.125 – 0.131) | 0.129<br>(0.127 – 0.168) | 0.517 |

\*Kruskal – Wallis Test

*Kruskal – Wallis Test

† IQR – Inter quartile Range, Q1 – 25^th^ Percentile, Q2 – 75^th^ Percentile

‡ WS – *Withania somnifera*

**Figure 1:**
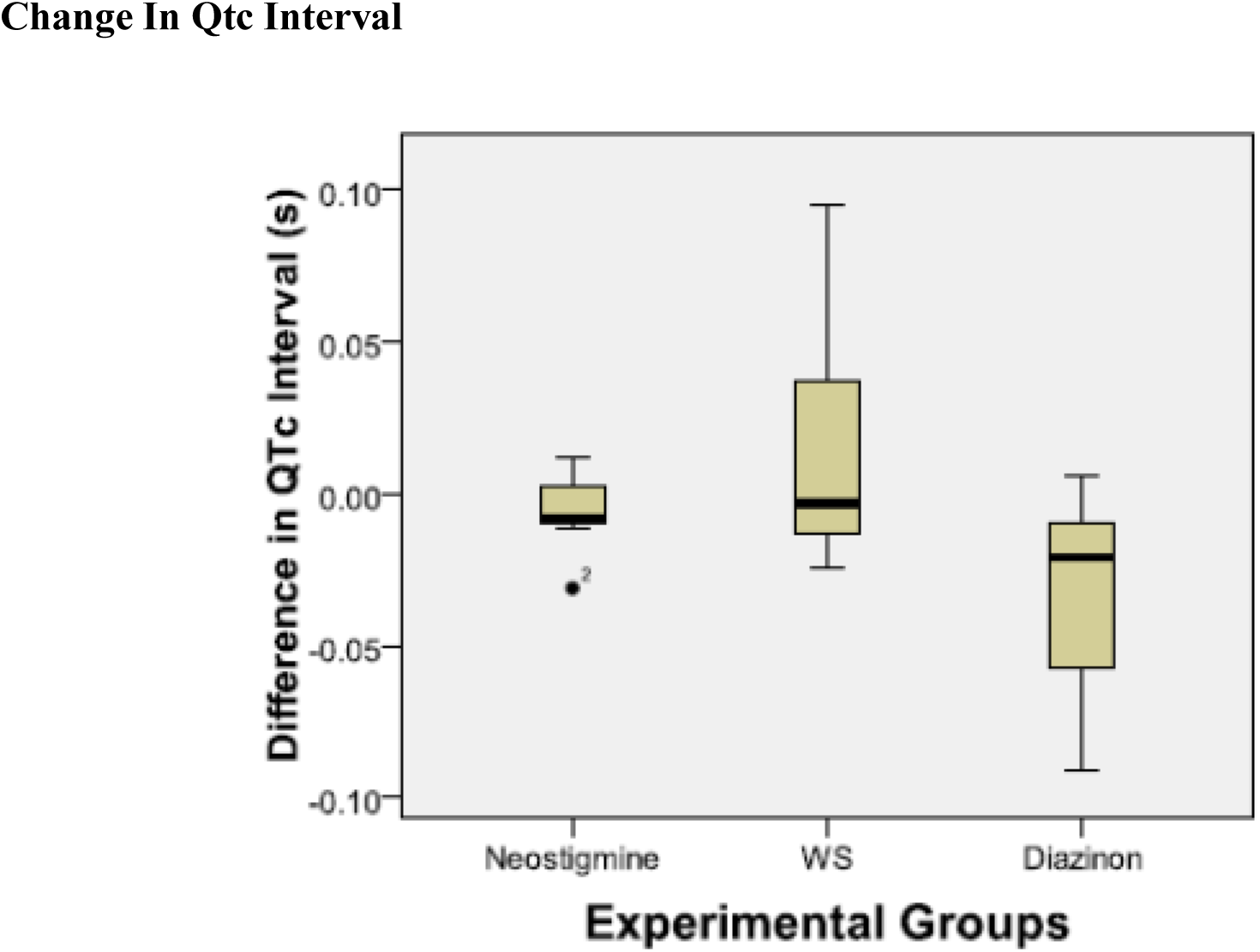
Box plots showing change in QTc interval (s) (post test - baseline).

### 3.1. RR Interval and Heart Rate

The change in RR intervals between the groups was statistically significant (P = 0.031) (Figure 2). However, Post hoc analysis showed no statistically significant difference in the change in RR interval between the positive control and test groups (0.026 (0.007 – 0.065) vs. 0.035 (0.019 – 0.050), P = 0.710). Accordingly, there was no significant change in the HR between the positive control and test groups (-54.235 (-115.317 – (-19.857)) vs. -−96.136 (-96.472 – (-43.879)), P = 0.902).

**Figure 2:**
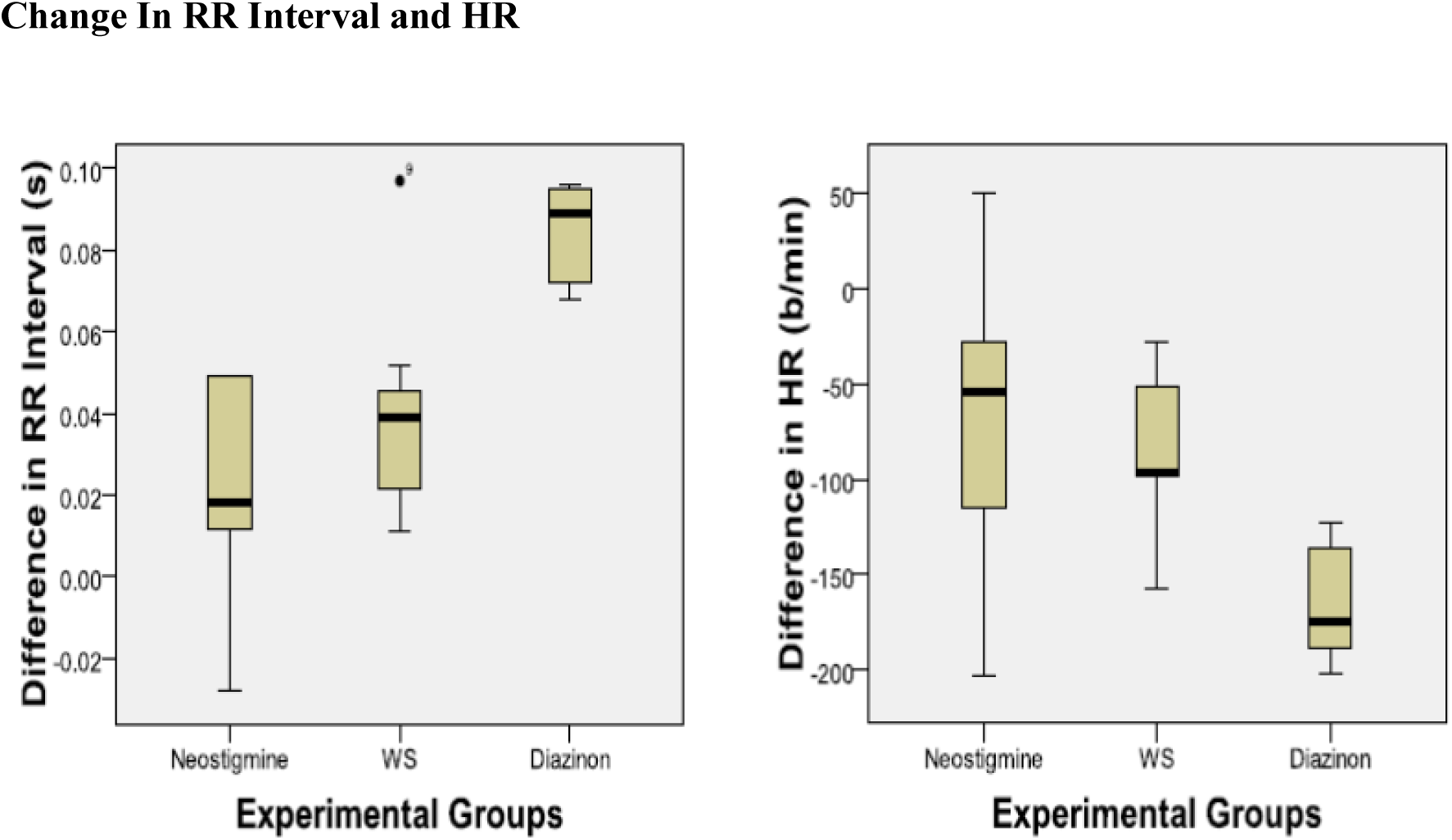
Box plots showing change in RR interval (s) and HR (b/min) (post test - baseline)

### 3.2. PR Interval

The change in PR intervals between the groups was statistically significant (P = 0.019) (Figure 3). The PR interval was significantly less shortened in the test group compared to the positive control group (0.003 (0.001 – 0.004) vs. 0.006 (0.004 – 0.008), (P = 0.038). Conversely, there was no statistically significant difference in the change in the PR interval between the positive control and negative control groups (0.006 (0.004 – 0.008) vs. 0.009 (0.008 – 0.015), P = 0.073).

**Figure 3:**
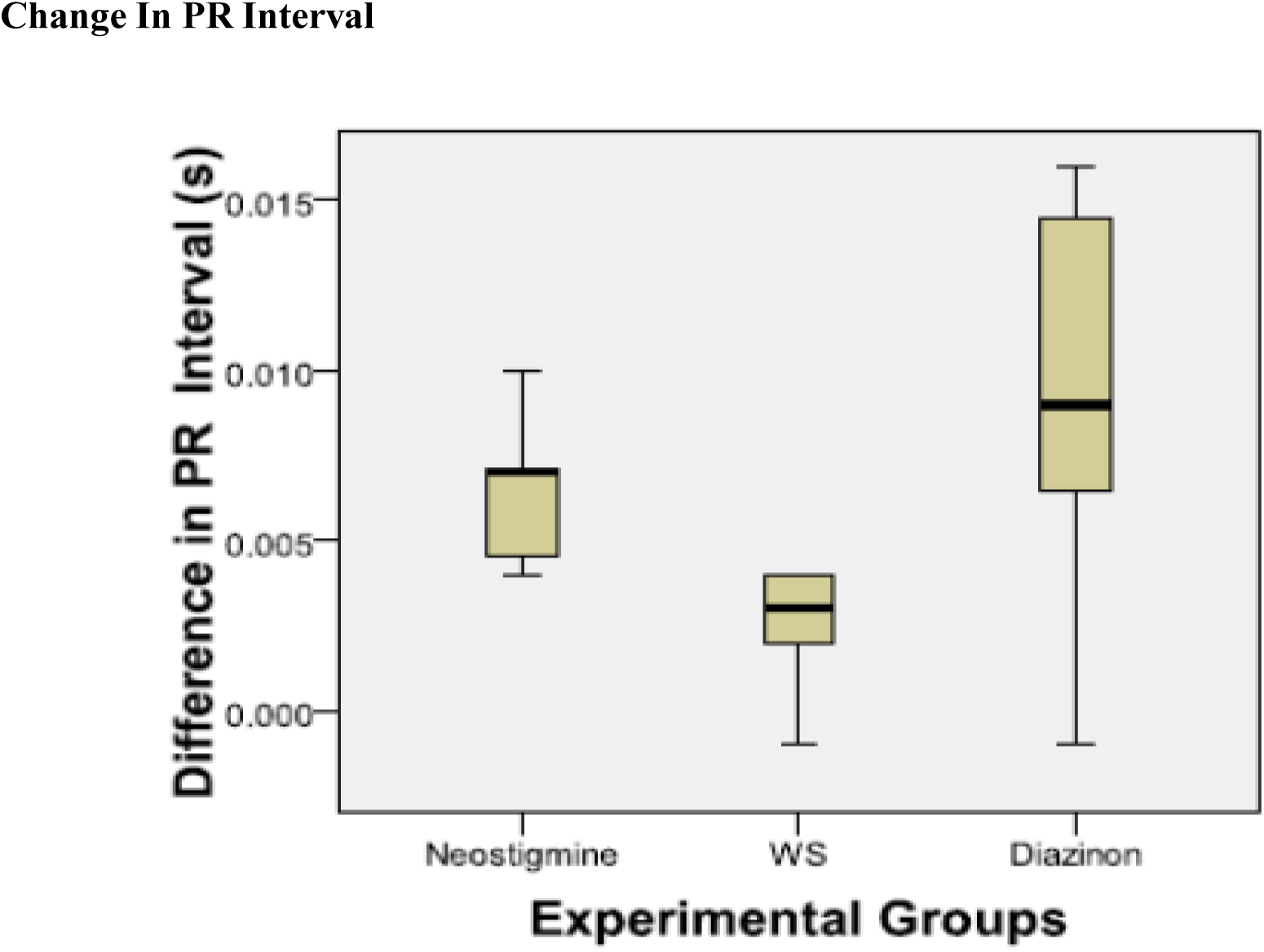
Box plots showing change in PR interval (s) (post test - baseline)

### 3.3. QRS Interval

There was a statistically significant difference in the change in QRS intervals between the groups (P = 0.023) (Figure 4). The Post hoc analysis however showed that there was no significant difference between the positive control and test groups (0.005 (0.001 – 0.007) vs. -−0.002 (-0.005 – 0.001), (P = 0.128).

**Figure 4:**
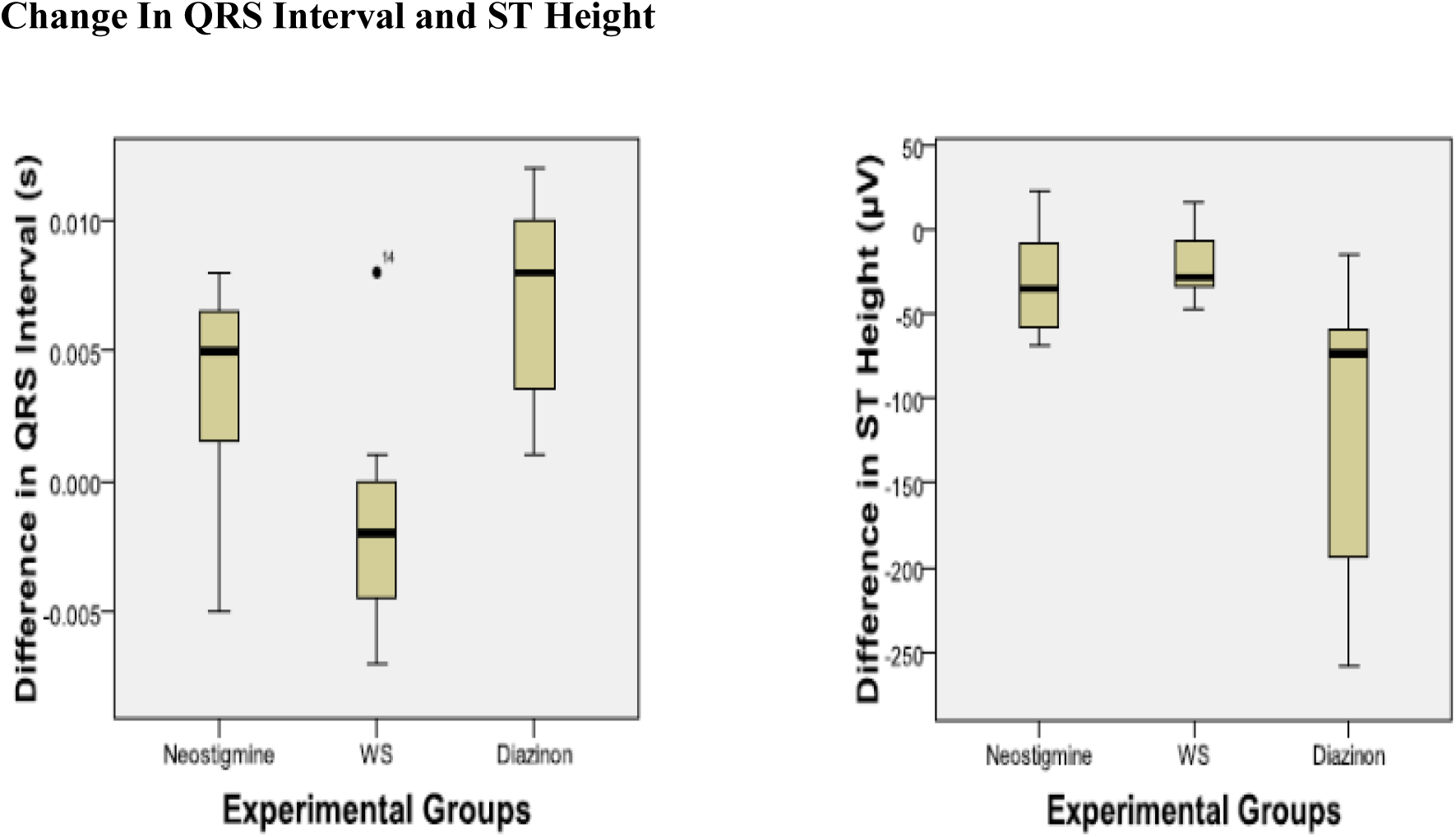
Box plots showing change in QRS interval (s) and ST Height (μV) (post test-baseline)

### 3.4. ST Height

There was a statistically significant difference in the change in ST height between the groups (P = 0.023) (Figure 4). The Post hoc analysis however showed that there was no significant difference between the positive control and test groups (-34.830 (-63.578 – 4.215) vs. -−22.330 (-38.383– (-4.159)), P = 0.535).

## 4.0. Discussion

Changes in the ECG are now recognized as among the first manifestations of cardiotoxicity and predate changes in the mechanical function of the heart (Gharib and Burnett, 2002). This is because the ECG changes are a reflection of the instantaneous changes in biochemical and metabolic processes that underlie cardiac function. The ECG therefore provides a convenient and rapid measure albeit indirect of the structural changes in the heart that will ultimately affect electromechanical function (Gharib and Burnett, 2002).

In common with other types of chemical cardiotoxicity, organophosphate poisoning is often accompanied by an arrhythmogenic phenotype having three well recognized stages. The initial phase is denoted by sinus tachycardia and hypertension due to sympathetic stimulation. This is then followed by hypotension, sinus bradycardia and artrio-ventricular node conduction disturbances associated with parasympathetic potentiation. The final stage is characterized by prolonged QT that often deteriorates into Torsades de Pointes and ventricular fibrillation which are often fatal (Abraham et al., 2001; Rubinstein et al., 1964). The cardiotoxic effects of organophosphates are the main cause of death in poisoning and are therefore a rational target for therapeutic intervention. This is especially in view of the fact that the Torsades de Pointes and ventricular fibrillation may suddenly develop up to fifteen days after exposure leading to sudden cardiac death after all the other signs of clinical intoxication have subsided (Bar-Meir et al., 2007;Viajakumar et al., 2011).

The exact mechanisms underlying cardiac toxicity are yet to be fully understood (Anand et al., 2009; Saadeh et al., 1997). However, it is well accepted that Organophosphate toxicity arises mainly due to acetylcholinesterase inhibition, which then leads to the accumulation of acetylcholine at cholinergic synapses (Namba, 1971). Therapeutic approaches to Organophosphate poisoning may be classified into antidotal post exposure treatments that aim to ameliorate the effects of cholinergic excess (Tenberken et al., 2010) and pre-treatment before exposure with subsequent antidotal treatment (Bajgar, 2004). The goal of pre-treatment is to keep synaptic acetylcholinesterase active by either protecting it from inhibition or by reducing the concentration of the Organophosphate at the synapse.

Much modern research on therapeutic management of Organophosphate poisoning has focused on the development of novel pre-exposure treatment regimens in view of its utility for use by categories of workers who have unavoidable exposure to Organophosphates, for instance, farm workers, military personnel, among others (Bajgar, 2004). These new protocols however are often expensive to administer and are consequently impractical in resource poor settings highlighting the need to develop newer, cheaper, efficacious and more effective treatment options for Organophosphate poisoning (Buckley et al., 1996). The objective of our study therefore was to evaluate the use of *Withania somnifera* as a low cost alternative to Neostigmine pre-treatment in the management of Organophosphate poisoning.

*Withania somnifera* is a perennial herb species that finds extensive application in Ayurvedic and African traditional medicine for the treatment of a wide variety of conditions (Singh & Kumar, 1998). It possesses reversible cholinesterase inhibitory activity similar to that of the carbamates (Choudhary et al., 2004). This anti-cholinesterase activity has been attributed to the withanolides, which are some of its major chemical constituents (Choudhary et al., 2004). Withanolides (chemical name: 22-hydroxy ergostane-26-oic acid 26, 22-d-lactones) are a group of naturally occurring, twenty eight (28) carbon steroids built on an ergostane framework with a lactone-containing side chain of nine carbons attached at the carbon seventeen (C-17) (Mirjalili et al., 2009) (Figure 5).

**Figure 5:**
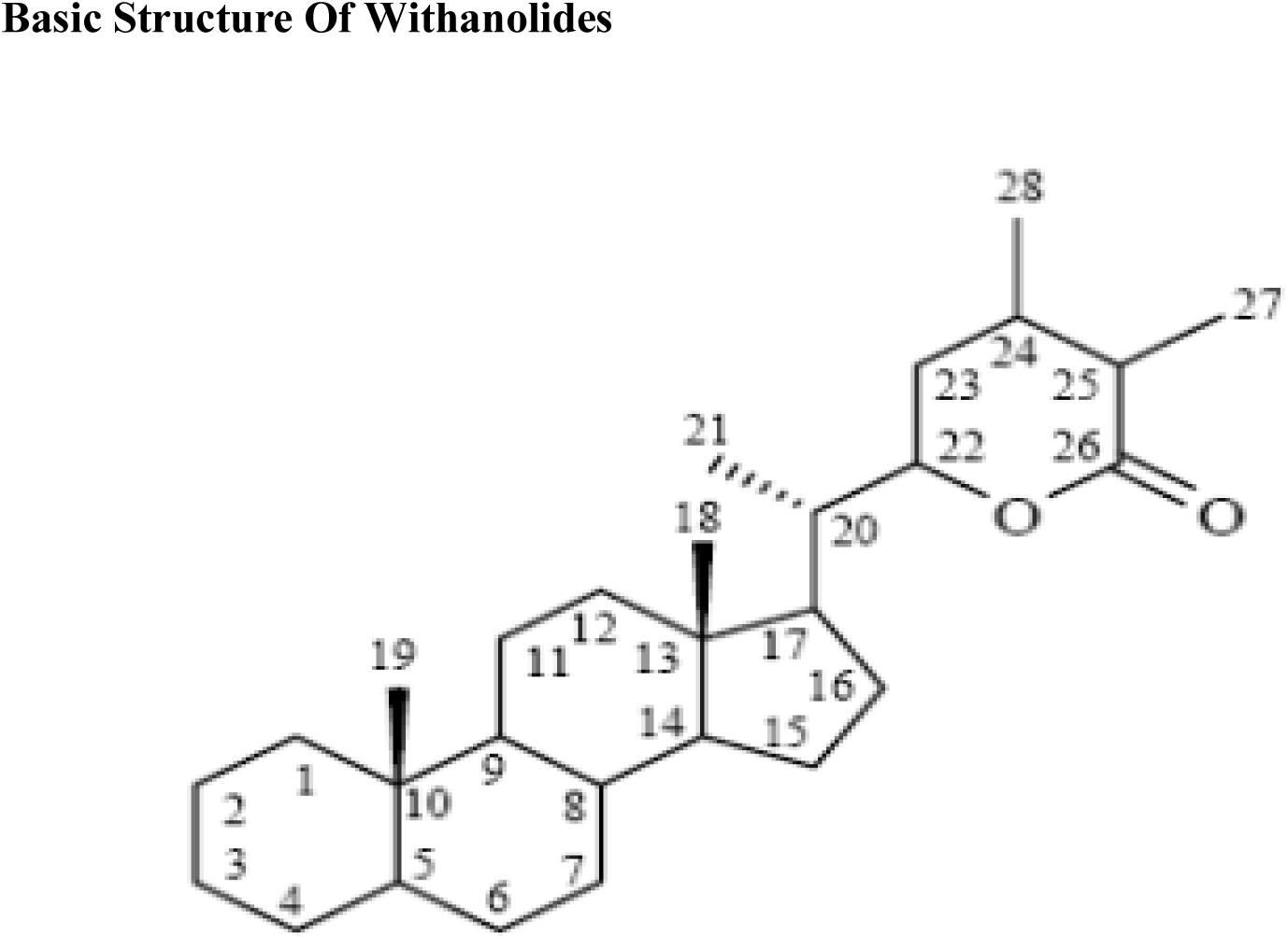
Basic structure of withanolides (Mirjalil et al. 2009, p. 2375)

Pre-treatment with the methanol extract of *Withania somnifera* had a significant effect in preventing changes in the following ECG parameters; RR interval, HR, PR interval, QRS interval and the ST segment. Even more importantly, there were no significant differences between pre-treatment with Neostigmine and pre-treatment with the methanol extract of *Withania somnifera*. This implies that there is no significant difference in efficacy between the extract and neostigmine.

Neostigmine is a carbamate drug that transiently carbamylates the active site of acetylcholinesterase causing reversible inhibition of the enzyme (Masson, 2011). Since it spontaneously dissociates from the enzyme, it helps to protect the active site from permanent inhibition by the Organophosphates and hence allows the preservation of a pool of protected enzyme that decarbamylates spontaneously to regenerate the acetlycholinesterase activity (van Helden et al., 2011). Hence the carbamates, with pyridostigmine (which is closely related to Neostigmine) as a salutary example, are used in pre-treatment drug regimens to prevent Organophosphate poisoning in various situations including wartime (Masson, 2011; Van Helden et al., 2011).

The carbamates however cause as side effects the spectrum of symptoms called the Gulf syndrome characterized by weakness, diarrhoea, nausea, vomiting, hyper peristalsis, excessive salivation, difficulty in breathing and development of CNS signs such as restlessness, dizziness and hallucinations (Weinbroum, 2004). These side effects make it mandatory to have medical staff in close attendance during and after their administration, a major drawback in resource limited settings. There are as yet no published reports of similar side effects being associated with the use of *Withania somnifera* despite having reversible acetylcholinesterase activity and this may be a distinct advantage associated with the use of this herb vis a vis the carbamates.

## 6.0. Conclusions

The results indicate that the methanol extract of *Withania somnifera* may be a viable pre-treatment alternative to the carbamates in particular Neostigmine. It is therefore a potentially valuable addition to the therapeutic armoury available to health workers in resource limited settings for the management of Organophosphate poisoning. Future studies should focus on trying to isolate the chemical moiety (-ies) responsible for the protective effect against the cardiotoxicity. The studies should also be repeated in animal species with higher predictive validity as far as application to man is concerned e.g. Guinea pigs and pigs.

